# Behavior can be decoded across the cortex when individual differences are considered

**DOI:** 10.1101/2024.03.12.584674

**Authors:** Johan Nakuci, Jiwon Yeon, Ji-Hyun Kim, Sung-Phil Kim, Dobromir Rahnev

## Abstract

Group-level analyses have typically linked behavioral signatures to a constrained set of brain areas. Here we show that two behavioral metrics – reaction time (RT) and confidence – can be decoded across the cortex when each individual is considered separately. Subjects (N=50) completed a perceptual decision-making task with confidence. We built models decoding trial-level RT and confidence separately for each subject using the activation patterns in one brain area at a time after splitting the entire cortex into 200 regions of interest (ROIs). First, we developed a simple test to determine the robustness of decoding performance, which showed that several hundred trials per subject are required for robust decoding. We then examined the decoding performance at the group and subject levels. At the group level, we replicated previous results by showing that both RT and confidence could be decoded from a small number of ROIs (12.0% and 3.5%, respectively). Critically, at the subject level, both RT and confidence could be decoded from most brain regions even after Bonferroni correction (90.0% and 72.5%, respectively). Surprisingly, we observed that many brain regions exhibited opposite brain-behavior relationships across individuals, such that, for example, higher activations predicted fast RTs in some subjects but slow RTs in others. All results were replicated in a second dataset. These findings show that behavioral signatures can be decoded from a much broader range of cortical areas than previously recognized and suggest the need to study the brain-behavior relationship at both the group and subject levels.

## Introduction

An enduring goal of cognitive neuroscience is to establish the relationship between brain and behavior. For example, studies have linked brain measures to healthy aging (Dosenbach et al., 2010; Geerligs et al., 2015), personality (Yarkoni, 2015), intelligence (Finn et al., 2015; Heuvel et al., 2009), and mood (Smith et al., 2015). Additionally, there are considerable efforts to use brain measures for diagnosis (Arbabshirani et al., 2013; Yahata et al., 2016), treatment selection (Williams et al., 2015), and prediction of patient outcomes in clinical settings (Whelan et al., 2014).

What is the right level at which to study the brain-behavior relationship? It is increasingly evident that this relationship is a complex interplay between group-level factors, which are shared among individuals, and individual-level factors that manifest as unique characteristics (Dubois and Adolphs, 2016; Gratton et al., 2018; Nakuci et al., 2023b). The discernible variation in brain activity among individuals has long been recognized for its potential to reveal the intricacies of the differences in behavior between individuals (Miller et al., 2012; van Horn et al., 2008). Understanding how the individual differs from the group is critical since individual factors may be crucial for diagnosing and treating pathology (Gratton et al., 2020; Lebreton et al., 2019). Consequently, it is imperative to understand the complex individual factors shaping the brain’s role in behavior.

Group-level analyses using both mass univariate statistics or multivariate pattern analysis with searchlight have been used to identify brain behavior signatures across the cortex (Friston et al., 1999; Kriegeskorte et al., 2008). These analyses typically uncovered behavioral signatures within constrained sets of brain areas. However, in the presence of substantial individual variability, one may expect that many areas of the brain that are not predictive of behavior in the group may nonetheless predict behavior in certain individuals. Nevertheless, it remains unknown how widely across cortex one can decode behavioral signatures in individual subjects. However, the problem is partly statistical since one requires both a lot of power (e.g., high number of trials) within an individual and a large enough sample of individuals for individual differences to manifest.

Differences across individuals manifest themselves in multiple ways. Previous work has found differences in spatial organization among brain regions across individuals (Williams et al., 2015). Moreover, there are substantial differences in large-scale organization among individuals (Braga and Buckner, 2017; Dworetsky et al., 2024). Crucially, these differences, particularly in spatial organization among individuals culminate in decoding across individuals being generally much poorer than within an individual (Haxby et al., 2011).

Here we investigate the extent to which two behavioral signatures – reaction time (RT) and confidence – can be decoded from across the cortex. Each subject (N=50) completed 700 trials from a perceptual decision-making task – a number significantly surpassing standard practice. The high number of trials empowers us to robustly estimate the brain-behavior relationship within individuals. To anticipate, we find that when factoring in individual differences, RT and confidence can be decoded from brain activity across the cortex in stark contrast to group-level analyses. We replicated these results in a second dataset where subjects (N=36) completed 804 trials of a different perceptual decision-making task. These results demonstrate that behavior can be predicted from a wider set of brain areas than would be suggested by standard group analyses.

## Methods

### Dataset 1 subjects and task

Dataset 1 has been previously published (Nakuci et al., 2023b). All details can be found in the original publication. Briefly, subjects (N= 50; 25 females; mean age = 26; age range = 19-40) completed a task where they judged which set of colored dots (red vs. blue) is more frequent in a cloud of dots. The stimulus was presented for 500 ms and subjects made untimed decision and confidence decisions using separate button presses. Most subjects performed six runs of 128 trials (a total of 768 trials). Three subjects completed only half of the 6^th^ run and another three subjects completed only the first 5 runs due to time constraints. All subjects were screened for neurological disorders and MRI contraindications. The study was approved by Ulsan National Institute of Science and Technology Review Board and all subjects gave written consent.

### Dataset 1 MRI recording

The MRI data was collected on a 64-channel head coil 3T MRI system (Magnetom Prisma; Siemens). Whole-brain functional data were acquired using a T2*-weighted multi-band accelerated imaging (FoV = 200 mm; TR = 2000 ms; TE = 35 ms; multiband acceleration factor = 3; in-plane acceleration factor = 2; 72 interleaved slices; flip angle = 90°; voxel size = 2.0 x 2.0 x 2.0 mm^3^). High-resolution anatomical MP-RAGE data were acquired using T1-weighted imaging (FoV = 256 mm; TR = 2300 ms; TE = 2.28 ms; 192 slices; flip angle = 8°; voxel size = 1.0 x 1.0 x 1.0 mm^3^).

### Dataset 2 subjects and task

Dataset 2 has been previously published as Experiment 2 in Yeon et al., 2020. All details can be found in the original publication. Briefly, subjects (N = 36, 23 females; mean age = 21.5; age range = 18-28) completed a task where they indicated whether a moving-dots stimulus had an overall coherent motion (always in downward direction) or not. The stimulus was presented for 500 ms and subjects made untimed decision immediately after the stimulus. All subjects completed 6 runs of 144 trials (a total of 864 trials). In the first half of the experiment (runs 1-3), subjects performed the task without providing confidence ratings. In the second half of the experiment (runs 4-6), subjects reported their confidence level with a separate, untimed button press immediately after making their perceptual decision. All subjects were screened for neurological disorders and MRI contraindications. The study was approved by the Georgia Tech Institutional Review Board and all subjects gave written consent.

### Dataset 2 MRI recording

The MRI data were collected on 3 T MRI systems using a 32-channel head coils. Anatomical images were acquired using T1-weighted sequences with a MEMPRAGE sequence, FoV□=□256 mm; TR□=□2530 ms; TE□=□1.69 ms; 176 slices; flip angle□=□7°; voxel size□=□1.0□×□1.0□×□1.0 mm^3^. Functional images were acquired using T2*-weighted gradient echo-planar imaging sequences (FoV□=□220 mm; TR□=□1200 ms; TE□=□30 ms; 51 slices; flip angle□=□65°; voxel size□=□2.5□×□2.5□×□2.5 mm^3^).

### MRI preprocessing

MRI data were preprocessed with SPM12 (Wellcome Department of Imaging Neuroscience, London, UK). All images were first converted from DICOM to NIFTI and removed the first three volumes to allow for scanner equilibration. We then preprocessed with the following steps: de-spiking, slice-timing correction, realignment, segmentation, coregistration, normalization, and spatial smoothing with 10 mm full width half maximum. Despiking was done using the 3dDespike function in AFNI. The preprocessing of the T1-weighted structural images involved skull-removal, normalization into MNI anatomical standard space, and segmentation into gray matter, white matter, and cerebral spinal fluid, soft tissues, and air and background.

### Single-trial beta estimation

Single-trial beta responses were estimated with a general linear model (GLM) using GLMsingle, a Matlab toolbox for single-trial analyses (Prince et al., 2022). The hemodynamic response function was estimated for each voxel and nuisance regressors were estimated in the same manner as described in Allen *et al*. (Allen et al., 2022). Additionally, regressors for the global signal and for six motion parameters (three translation and three rotation) were included. The single-trial betas were estimated in three batches. In each batch, the betas for every third trial were estimated because the trials in our study were temporally close together. Also, trials that were within 20 seconds from the end of run were removed due to the lack of sufficient length of signal from which to estimate the trial-specific hemodynamic response function. The beta estimates from GLMsingle have been validated in our previous work and the results are comparable to what standard GLM analyses is SPM12 (Nakuci et al., 2023a). Further, to reduce any differences in the modeling training and fitting that might arise from difference in the number of trials between subject, when possible, we opted for a uniform number of trials per subject. In total, the analysis on was based on 700 trials for Dataset 1 and 804 trials for Dataset 2; for subjects who had more trials than these, we simply removed trials from the end of the experiment. (Note that in six subjects in Dataset 1 and two subjects in Dataset 2 had fewer total trials because they did not complete all 6 runs.)

### Subject-level brain-behavior decoding analysis

For each trial, the activation within each of the 200 regions of interest (ROIs) from the Schaefer atlas was estimated by averaging all voxel in the ROI (Schaefer et al., 2018). Individual trials were randomly separated into training and testing bins each containing 350 trials in Dataset 1 and 402 trials in Dataset 2. (For the confidence analyses in Dataset 2, the training was done on 301 trials and testing on the remaining 101 trials.) For each ROI, a linear model was trained on training bin and used to predict behavioral performance on the testing bin. The linear model was trained using *fitlm.m* in Matlab. Additionally, we utilized a more advanced model based on Support Vector Regression (SVR) to determine if decoding could be improved when compared to the simple linear model. The SVR model was trained using *fitrsvm.m* in Matlab with default parameters.

Decoding performance for a given ROI was determined by correlating the empirical and predicted RT and confidence. The analysis was repeated 25x to ensure that model performance was not dependent on the initial division of trials. The decoding performance values in each of the 25 iterations were first z-scored using the Fisher transformation and then averaged and converted back to r-values. Significance was determined by converting r-values to t-values.

Decoding analysis was conducted for each individual subject separately and compared to a permuted null model. A brain region was deemed to significantly decode behavioral performance if the correlation between predicted and empirical RT or confidence exceeded a null model based on permutating RT or confidence 1000x at P < 0.05, uncorrected for multiple comparison (P_uncor_).

However, when conducting an analysis at the individual level, multiple comparisons is a particularly acute issue because comparisons are performed independently across many subjects (50 in Experiment 1 and 36 in Experiment 2) and across 200 regions within a subject. Therefore, a multiple comparison correction is needed. In Dataset 1, subject-level decoding performance was evaluated at P < 0.05 uncorrected and with three Bonferroni multiple comparison corrections of 50 tests (equal to the number of subjects), 200 tests (equal to the number of ROIs), and 10,000 tests (the number of subjects times the number of ROIs). Similarly, in Dataset 2, subject-level decoding performance was evaluated at P < 0.05 uncorrected and with three Bonferroni multiple comparison corrections of 36 tests (equal to the number of subjects), 200 tests (equal to the number of ROIs), and 7,200 tests (the number of subjects times the number of ROIs).

### Group-level brain-behavior decoding analysis

Group-level decoding performance was estimated by averaging the individual subject decoding performance for each ROI. Group-level decoding was Bonferroni corrected for 200 tests (equal to the number of ROIs).

### Decoding performance and trial number

We developed a test to determine how many trials are needed to obtain robust individual-level brain-behavior relationships. The test relies on estimating decoding performance on a range of trials in training and testing bins. Specifically, for each subject, a whole brain multilinear regression model was trained on a subset of trials that ranged from 5% to 95% of trials and tested on the remaining trials. The multilinear regression model was repeated 25 times training on different subset of data. Decoding performance was estimated by averaging across the 25 iterations and decoding performance variance was estimated by calculating the standard deviation across the 25 iterations.

### Data and code

Data and scripts necessary to perform the analysis are available at https://osf.io/dc9pa/.

## Results

### Analytical framework for behavior decoding

We investigated the brain-behavior relationship in two perceptual decision-making datasets. In the first dataset, each subject (N=50) completed over 700 trials of a perceptual task with confidence. In the second dataset (N=36), each subject completed 804 trials but only half of the trials included confidence ratings. We recorded subjects’ reaction time (RT) and confidence and investigated how well they can be decoded from different parts of the brain. We utilized a recently developed method, GLMsingle, to estimate the voxel-wise activation on a given trial (i.e., single-trial beta) (Prince et al., 2022). For each subject, we performed the decoding analysis on the average activation within each of the 200 regions of interest (ROIs) from the Schaefer atlas (Schaefer et al., 2018). Specifically, for each trial, we averaged the beta values from all voxels within a brain region to obtain a single beta value for each ROI. Individual trials were then randomly separated into training and testing bins (**Fig. 1A**). A prediction model was trained on half of the trials (**Fig. 1B**, *left*) and used to predict behavioral performance on the remaining trials from the activation (beta values) separately for each ROI (**Fig. 1B**, *middle*). Decoding performance for a given ROI and subject was determined by correlating the empirical and predicted RT and confidence (**Fig. 1B**, *right*). The analysis was repeated 25x to ensure that model performance did not depend on the initial trial split (**Fig. 1C**).

**Figure 1.**
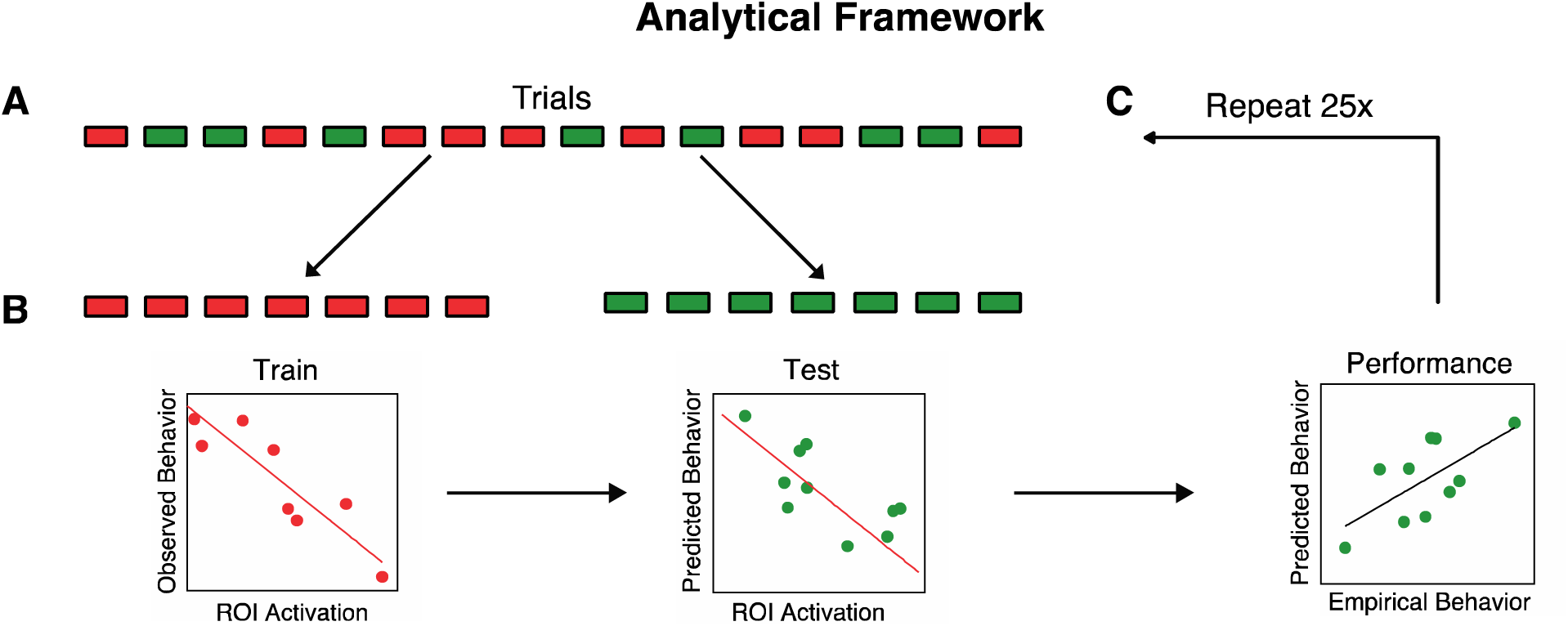
Analytical framework for behavior decoding analysis. (A) For each brain region, individual trials were randomly separated into training (red) and testing (green) bins. (B) The trials in the training bin were used to train a prediction model (*left)*. The model was used to predict the behavioral performance of the trials in the testing bin using the observed brain activation (*middle*). Model performance was estimated using the Pearson correlation between the empirical and predicted behavioral performance (*right*). (C) The analysis was repeated 25x to ensure that model performance was robust and did not depend on the initial trial split.

### Robust decoding performance requires several hundred trials per subject

Our Datasets 1 and 2 included 700 and 804 trials per subject, respectively. We first investigated how the number of trials used for training and testing affects decoding performance. We develop a simple test to determine how many trials are needed to obtain robust individual-level decoding.

The test relies on estimating decoding performance using a range of trials for both training and testing. For robustness, we used the data from all 200 ROIs to decode both RT and confidence. We trained a linear regression model on subsets of trials that ranged from 5% to 95% of all trials and tested on the remaining trials. For each percentage of trials used, we randomly selected trials for training and testing 25 times. We then computed both the average decoding performance across the 25 iterations and the variance in decoding performance. We then determined the percentage of trials that should be devoted to training the decoding model to minimize the variance in the decoding performance.

For Dataset 1, minimum decoding variance was obtained when training/testing each contained 350 of trials (50% of trials) for both RT (**Fig. 2A**) and confidence (**Fig. 2B**), while average decoding performance increased with higher number of training trials. Critically, using fewer than 350 trials for training exhibited poor decoding performance. On the other hand, using more than 350 trials for training improved performance, but the variance in performance increased across iterations. The maximum difference between decoding performance and variance was obtained when training was performed on 525 trials (75% of all trials). We found similar results for Dataset 2. Minimum decoding variance was obtained when training contained 402 trials (50% of trials) for RT (**Fig. 2C**) and 301 trials (75% of trials) for confidence (**Fig. 2D**), while decoding performance increased with higher number of training trials. These results suggest that a 50:50 split between training and testing bins might produce more consistent results compared to 80:20 or 90:10 (5- or 10-fold) division of trials between training and testing bins. All subsequent analyses are based on 50:50 split between training and testing bins which exhibited the optimal performance while minimizing the variance across iterations compared to other division ratios. Additionally, we repeated the analysis using a fixed number of trials during testing, 35 (5%), to equate performance across the different training sets. The result indicated that decreasing the number of trials used in training increased the decoding variance across iterations (**Fig. S1)**. Importantly, these results suggest the need of several hundred trials to robustly train a decoding model for RT or confidence, implying that many previous studies estimating brain-behavior relationship at the level of the individual may be underpowered.

**Figure 2.**
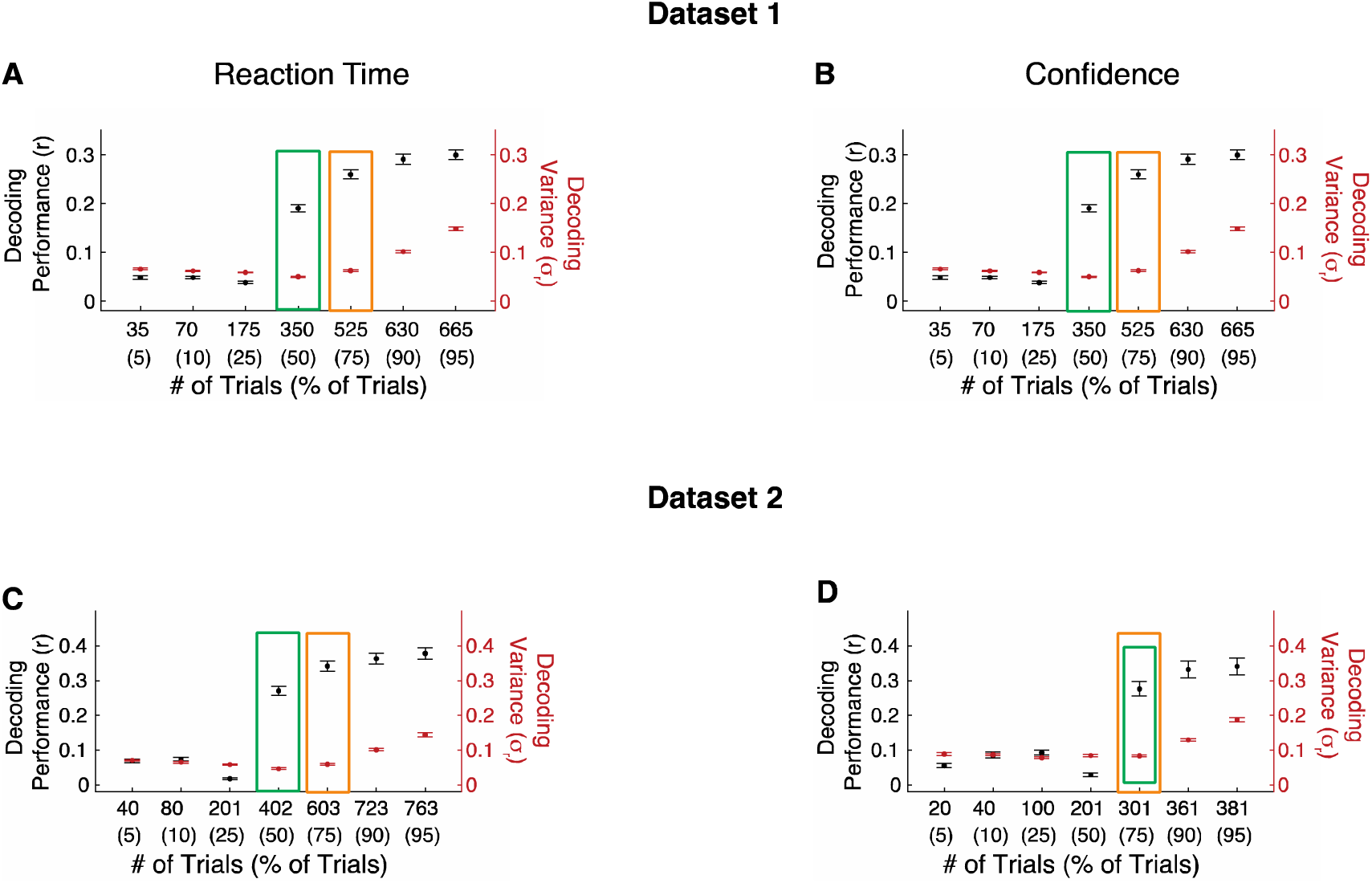
Robust decoding of RT and confidence requires several hundred trials. In Dataset 1, the variance in decoding performance was minimized when using 350 trials (corresponding to 50% of all trials) for both (A) RT and (B) confidence. The black dots present the average decoding performance across subject after 25 repeats for each subject (*left axis)*. The red dots present the average subject-level decoding variance across the 25 repeats for each subject (*right axis)*. The green boxes indicate the number of trials that minimize the variance in the decoding performance. The orange boxes indicate the number of trials that maximize the difference between decoding performance and the decoding variance. Error bars show SEM. (C-D) Same as Panels A-B but for Dataset 2. Note that in Dataset 2, confidence was measured on only have the trials (402) and correspondingly the decoding variance was minimized for a higher percentage of trials (75%).

### RT and confidence can be predicted from across the brain

Having examined how the number of trials used for training and testing affects decoding performance, we proceeded to investigate how widely across the brain we can decode RT and confidence. We first performed group-level analyses to decode RT across all subjects for each of the 200 ROIs. To identify brain regions from which RT and confidence could be significantly decoded at the group-level, the decoding performance values across individual subjects were aggregated and tested against zero. We found that RT could be significantly decoded in 12.0% of all ROIs after correcting for multiple comparisons (P < 0.05, Bonferroni-corrected, one-tailed one sample t-tests; **Fig. 3A**). These standard group-level analyses appear to suggest that RT can be decoded from only a handful of ROIs in the brain and that the rest of the ROIs cannot be used for decoding RT. We found similar results when we attempted to decode confidence instead of RT. At the group level, confidence could be decoded from only 3.5% of ROIs after correcting for 200 comparisons (one for each ROI; **Fig. 3B**). Moreover, we repeated our analysis using support vector regression model (SVR) and found similar results for both RT (**Fig. 2C**) and confidence (**Fig. 2D**) suggesting that more advanced models may only marginally improve decoding. The analyses focus on the results obtained with the linear model because of challenges associated with interpreting beta values from non-linear decoding models (Kriegeskorte et al., 2008).

**Figure 3.**
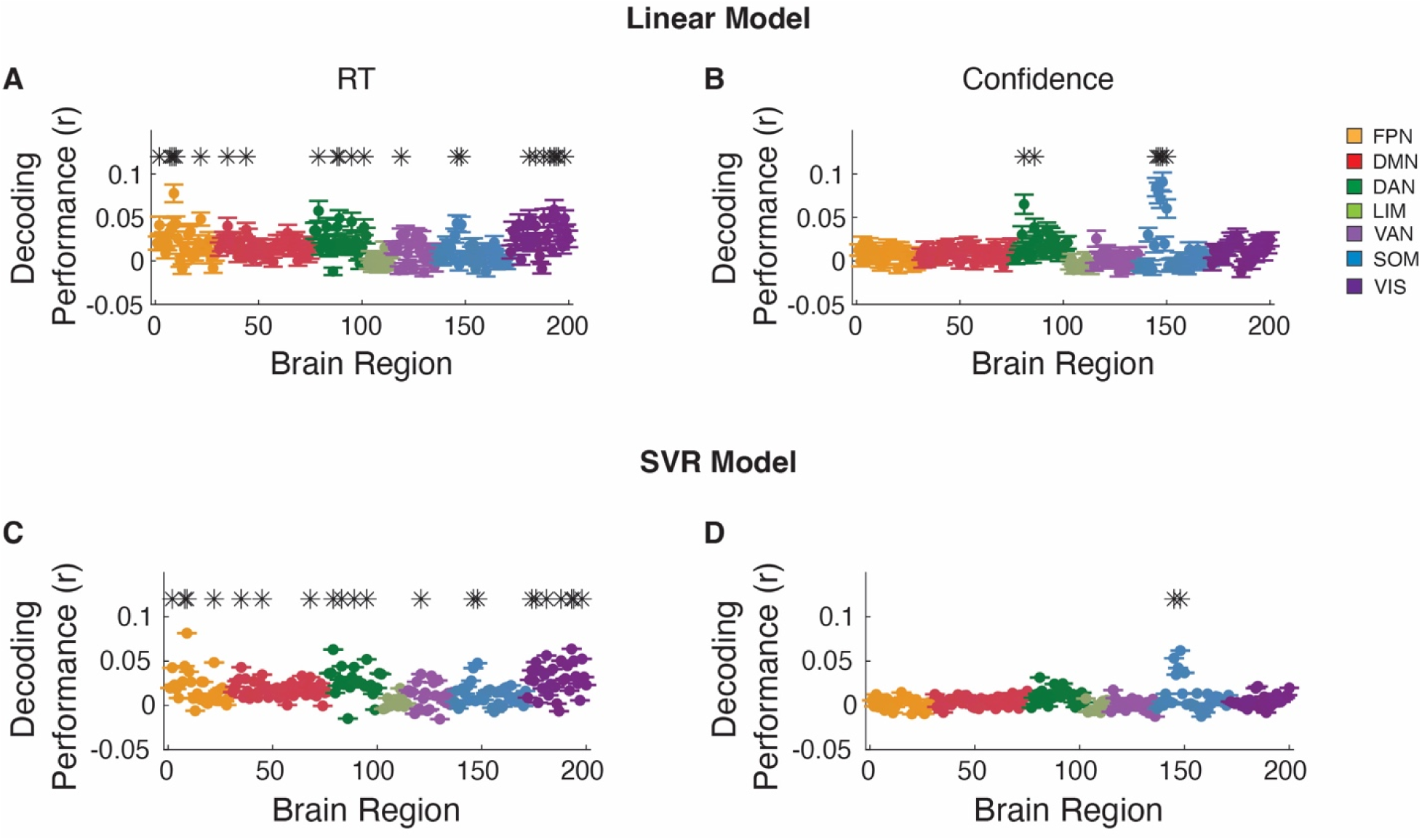
Group-level RT and confidence decoding in Dataset 1. Brain regions from which (A) RT and (B) confidence could be significantly decoded from at the group level after correcting for 200 comparisons. (C, D) Same as panel A and B, but using support vector regression (SVR) model to decode RT and confidence, respectively. * Indicates brain region that significantly, P < 0.05, Bonferroni corrected. FPN, Frontal Parietal Network; DMN, Default Mode Network; DAN, Dorsal Attention Network; LIM, Limbic Network; VAN, Ventral Attention Network; SOM, Somatomotor Network; VIS, Visual Network.

Critically, we asked whether this conclusion holds on the level of the individual subject. To address this question, we determined for how many ROIs there was at least one subject for whom RT could be decoded. We found that with no correction for multiple comparisons, decoding was significant for at least one subject in every single one of the 200 ROIs (**Fig. 4A-B**). Moreover, decoding performance at the individual level exceeded group-level performance. Additionally, as can be observed, there was substantial individual variability in decoding capabilities of ROIs across subjects (**Fig. 4B**).

**Figure 4.**
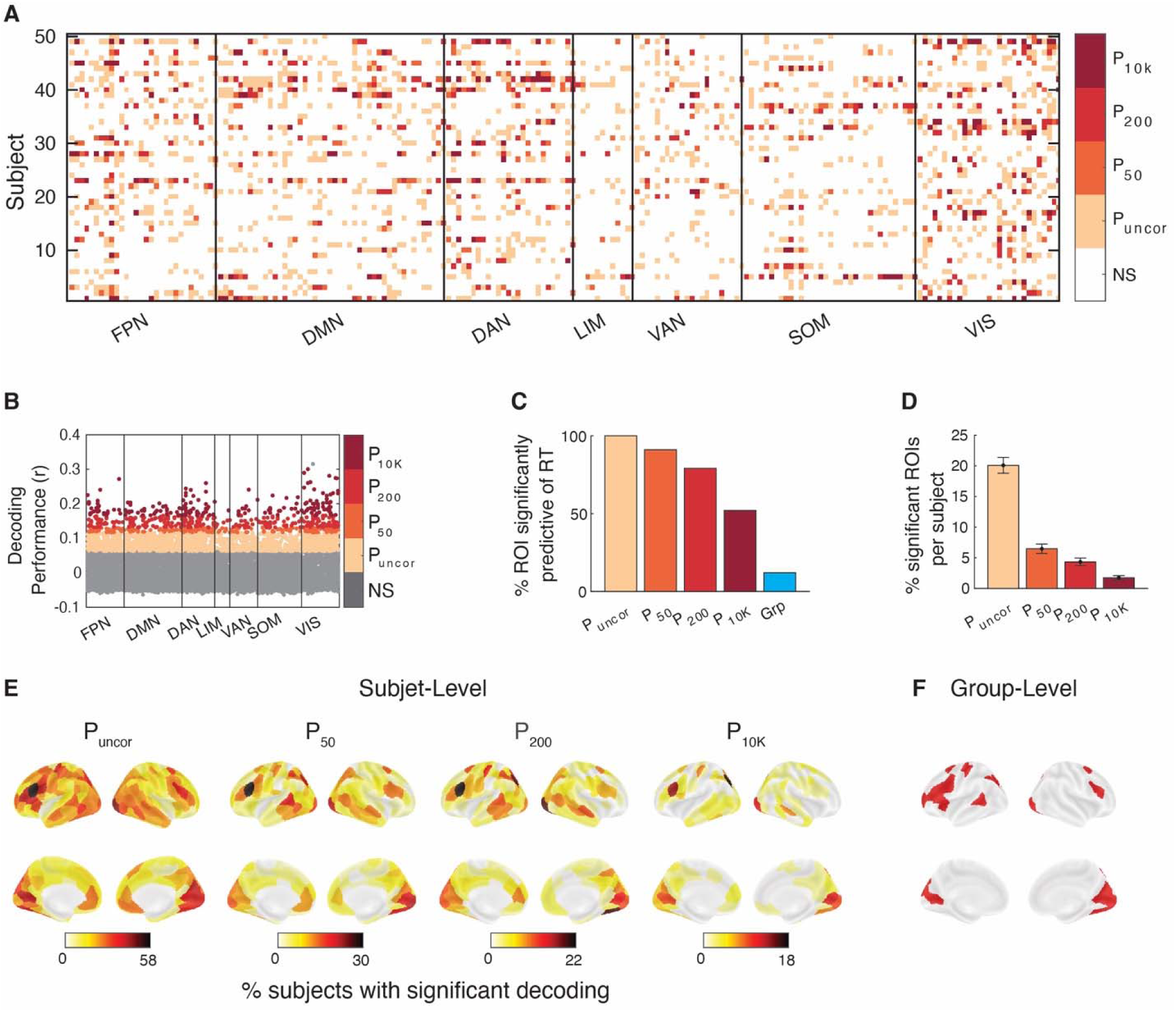
RT can be predicted from across the brain in Dataset 1. (A) Brain regions from which RT could be significantly decoded in each subject and region of interest. Significance is shown at four levels of correction: without any correction (P_uncor_), and after correcting for 50 (P_50_), 200 (P_200_), and 10,000 (P_10K_) comparisons. NS, not significant. Colors indicate the most conservative threshold at which one can significantly decode RT from a given region (see color legend on the right). (B) Decoding performance for which subject and brain region. Performance was estimated using the Pearson correlation between empirical and predicted RT. (C) Percentage of ROIs where RT could be significantly decoded for at least one subject for four levels of multiple comparison correction. Group-level results have been added for comparison. (D) Percentage of ROIs per subject where RT could be significantly decoded at four levels of multiple comparison correction. (E) Brain maps plotting percentage of subjects for whom RT could be significantly decoded for four levels of multiple comparison correction. (F) Brain maps plotting ROIs for which RT could be significantly decoded at the group-level after multiple comparison correction. FPN, Frontal Parietal Network; DMN, Default Mode Network; DAN, Dorsal Attention Network; LIM, Limbic Network; VAN, Ventral Attention Network; SOM, Somatomotor Network; VIS, Visual Network.

Next, we examined the decoding accuracy for each ROI separately and applied Bonferroni correction for the presence of 50 tests (equal to the number of subjects). We found that RT could be significantly decoded in at least one subject from 90% of all ROIs (180 of 200; **Fig. 4C**). These ROIs spanned all seven major brain networks associated with the Schaefer atlas. Further, when using a more stringent Bonferroni correction for 200 tests (equal to the number of ROIs), RT could be significantly decoded in at least one subject from 78.5% of all ROIs (157 of 200; **Fig. 4C**). Even with most the stringent Bonferroni correction for 10,000 tests (the number of subjects times the number of ROIs), RT could be significantly decoded in at least one subject from 50.5% of all ROIs (101 of 200, **Fig. 4C**). (Note, we show all four levels of correction because there is no “right” level of correction, and no level of correction communicates all relevant information.)

Importantly, even though most ROIs permitted the decoding of RT in at least one subject, this decoding was only significant in a few subjects per ROI. Specifically, the average ROI could be used to decode RT in 20.2% without multiple comparison correction, and in 6.6, 4.5, and 1.9% of subjects after correcting for 50, 200, and 10,000 comparisons (**Fig. 4D**). In all four cases, the most predictive ROIs were interspersed across much of the cortical surface compared to the group-level which was limited to a subset of regions (**Fig. 4E-F**). These results suggest the presence of strong individual differences, such that each ROI is only predictive of RT in a handful of subjects.

We found similar results when we attempted to decode confidence. Nevertheless, at the individual level, many more ROIs could be used to significantly decode confidence in at least one subject. Specifically, confidence could be decoded in 99.5% of all ROIs without multiple comparison correction, and in 72.5, 59.5, and 26.5% of ROIs after correcting for 50, 200, and 10,000 comparisons (**Fig. 5**). Note that confidence was overall less decodable than RT. Overall, the decoding results demonstrate that RT and confidence can be decoded from across the cortex when individual differences are considered.

**Figure 5.**
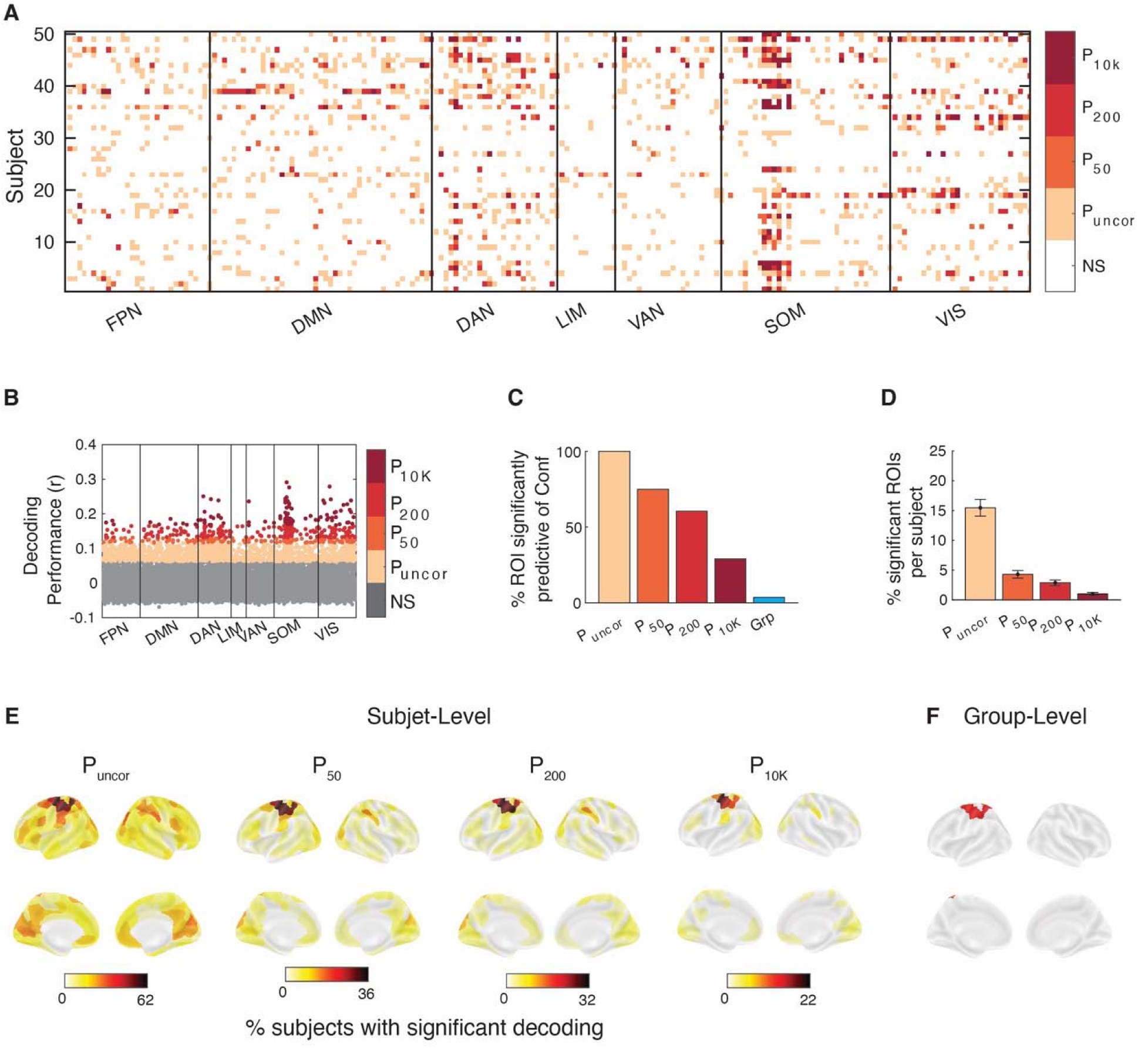
Confidence can be predicted from across the brain in Dataset 1. (A) Brain regions from which confidence could be significantly decoded in each subject and region of interest. Significance is shown at four levels of correction: without any correction (P_uncor_), and after correcting for 50 (P_50_), 200 (P_200_), and 10,000 (P_10K_) comparisons. NS, not significant. Colors indicate the most conservative threshold at which one can significantly decode confidence from a given region (see color legend on the right). (B) Decoding performance for which subject and brain region. Performance was estimated using the Pearson correlation between empirical and predicted confidence. (C) Percentage of ROIs where confidence could be significantly decoded for at least one subject for four levels of multiple comparison correction. Group-level results have been added for comparison. (D) Percentage of ROIs per subject where confidence could be significantly decoded at four levels of multiple comparison correction. (E) Brain maps plotting percentage of subjects for whom confidence could be significantly decoded for four levels of multiple comparison correction. (F) Brain maps plotting ROIs for which confidence could be significantly decoded at the group-level after multiple comparisons correction. FPN, Frontal Parietal Network; DMN, Default Mode Network; DAN, Dorsal Attention Network; LIM, Limbic Network; VAN, Ventral Attention Network; SOM, Somatomotor Network; VIS, Visual Network.

### Opposite brain-behavior relationship among subjects for the same brain region

Critically, for many ROIs, the relationship between brain activity and both RT and confidence often went in the opposite direction for different subjects. For example, one set of subjects would exhibit *higher* RT or confidence with higher ROI activation, whereas a different set of subjects would exhibit *lower* RT or confidence with higher ROI activation (**Fig. 6A-B**). Indeed, we found that without multiple comparison corrections, high brain activity predicted higher RT in at least one subject and lower RT in at least one subject in 67.0% of all ROIs (**Fig. 6C**). This percentage decreased to 16.5, 8.0, and 1.0% after correcting for 50, 200, and 10,000 comparisons, respectively. Interestingly, the dorsal attention network contained the most regions with opposite relationship between brain activity and RT among subjects, with the default mode and visual networks as close second and third, respectively (**Fig. 6D**). We found similar results for confidence with 75.5% of all ROIs showing significant decoding in different direction for at least two subjects, and 15.0, 7.5, and 0% after correcting for 50, 200, and 10,000 comparisons, respectively (**Fig. 6E-G**). The ROIs from which RT and confidence could be decoded at the group-level were the ROIs from which behavioral performance could be decoded in most subjects. These ROIs exhibited opposite brain-behavior relationship in 69.56% of subjects in RT and in 85.71% of subjects in confidence. Further, the default mode network contained the most regions with opposite relationship between brain activity and confidence among subjects (**Fig. 6H**). In contrast to the findings for RT that lacked large clusters where decoding was consistently in the same direction, we found that a subset of ROIs associated with somatomotor network exhibited consistent relationship between brain activity and confidence across subjects. Except for this relatively small cluster, our results demonstrate that the relationship between brain activity and behavioral outcomes is not universal and that it frequently goes in opposite directions for different subjects.

**Figure 6.**
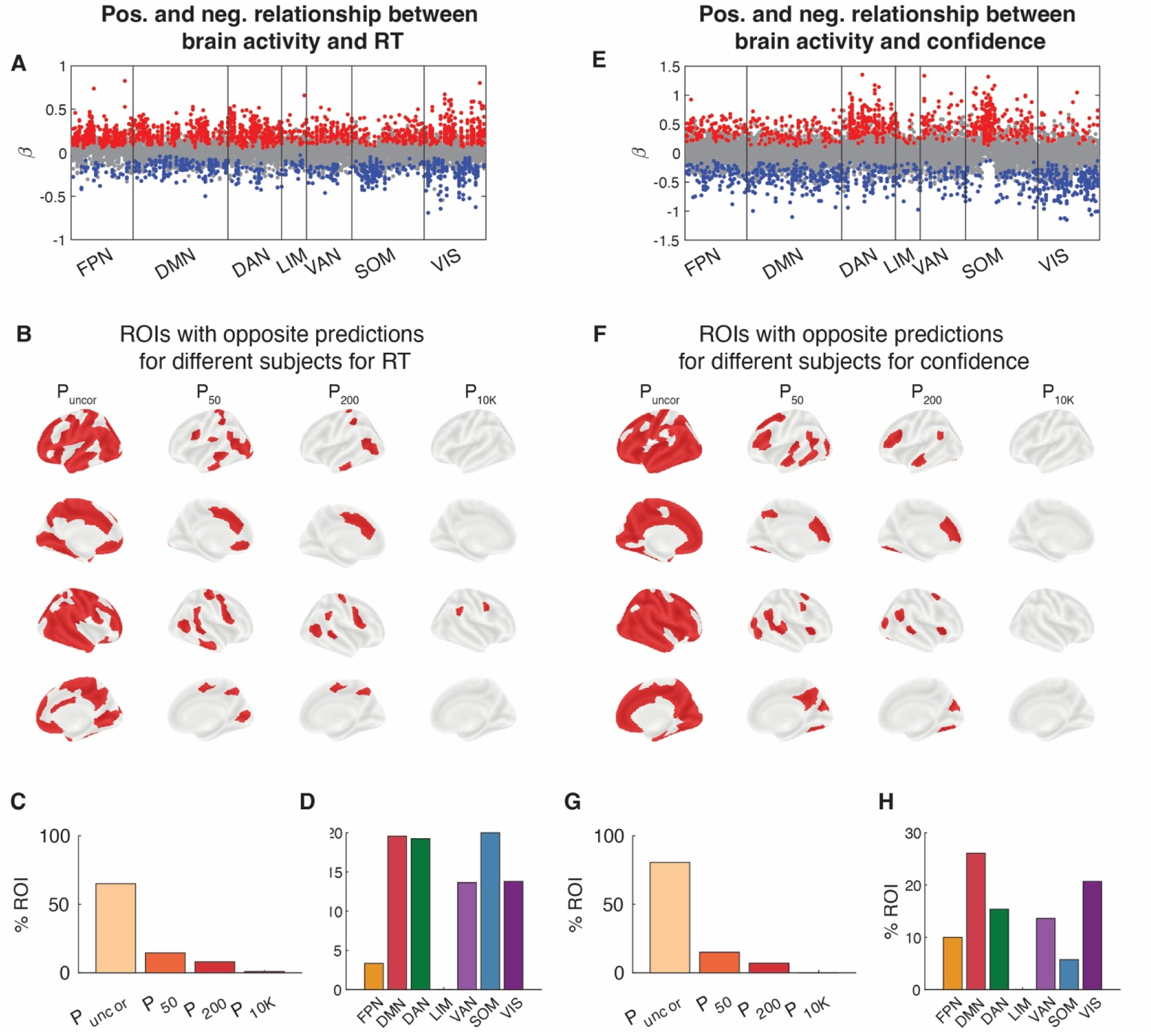
Opposite relationship between brain activity and both RT and confidence among subjects for the same brain region in Dataset 1. (A) Beta values for each subject and region of interest. Red and blue dots reflect subjects for whom RT could be significantly decoded from a given ROI (P < 0.05, uncorrected for multiple comparison). Gray dots show subjects for whom RT could not be significantly decoded from a given ROI. (B) Brain maps showing the ROIs that contained at least one subject with both positive and negative beta values for RT. Significance is shown at four levels of correction: without any correction (P_uncor_), and after correcting for 50 (P_50_), 200 (P_200_), and 10,000 (P_10K_) comparisons. (C) Percentage of ROIs that contained at least one subject with both positive and negative beta values for RT for each for four levels of multiple comparison correction. (D) Percentage of ROIs within each brain network that that contained at least one subject with both positive and negative beta values for RT after correcting for 50 comparisons. (E-H) Same as Panels A-D but for confidence.

### RT and confidence can be decoded from across the brain in Dataset 2

We replicated these results in a second dataset where subjects completed a different perceptual decision-making task with confidence (Yeon et al., 2020). Subjects (N=36) completed 804 trials but only half of them included confidence ratings, thus substantially decreasing the power for the confidence analyses.

Similar to Dataset 1, we found that at the group-level, RT could be significantly decoded in 13.0% of all ROIs after correcting for 200 comparisons in the linear model (P < 0.05, Bonferroni-corrected, one-tailed one sample t-tests; **Fig. 7A**). Crucially, we determined for how many ROIs there was at least one subject for whom RT could be decoded. At the individual level, RT could be decoded in 100% of all ROIs without multiple comparison correction, and in 87.0, 72, and 43.5% of ROIs after correcting for 36 (number of subjects), 200 (number of ROIs), and 7,200 (subjects times ROIs) comparisons (**Fig. 8**). Given the lower number of trials that contained confidence, we found that with the linear model, confidence could be significantly decoded in only 1.5% of all ROIs at the group-level after correcting for multiple comparisons (P < 0.05, Bonferroni-corrected, one-tailed one sample t-tests; **Fig. 7B**). Additionally, comparable results were obtained when using the SVR model for both RT and confidence (**Fig. 7C and D**). Despite the lower power, at the individual level, confidence could still be decoded in 76.5 % of all ROIs without multiple comparison correction, and in 47.0, 32.5, and 22.5% of ROIs after correcting for 36, 200, and 7,200 comparisons (**Fig. 9**). Overall, the results from Dataset 2 further support the notion that both the RT and confidence can be decoded from across the cortex when individual differences are considered.

**Figure 7.**
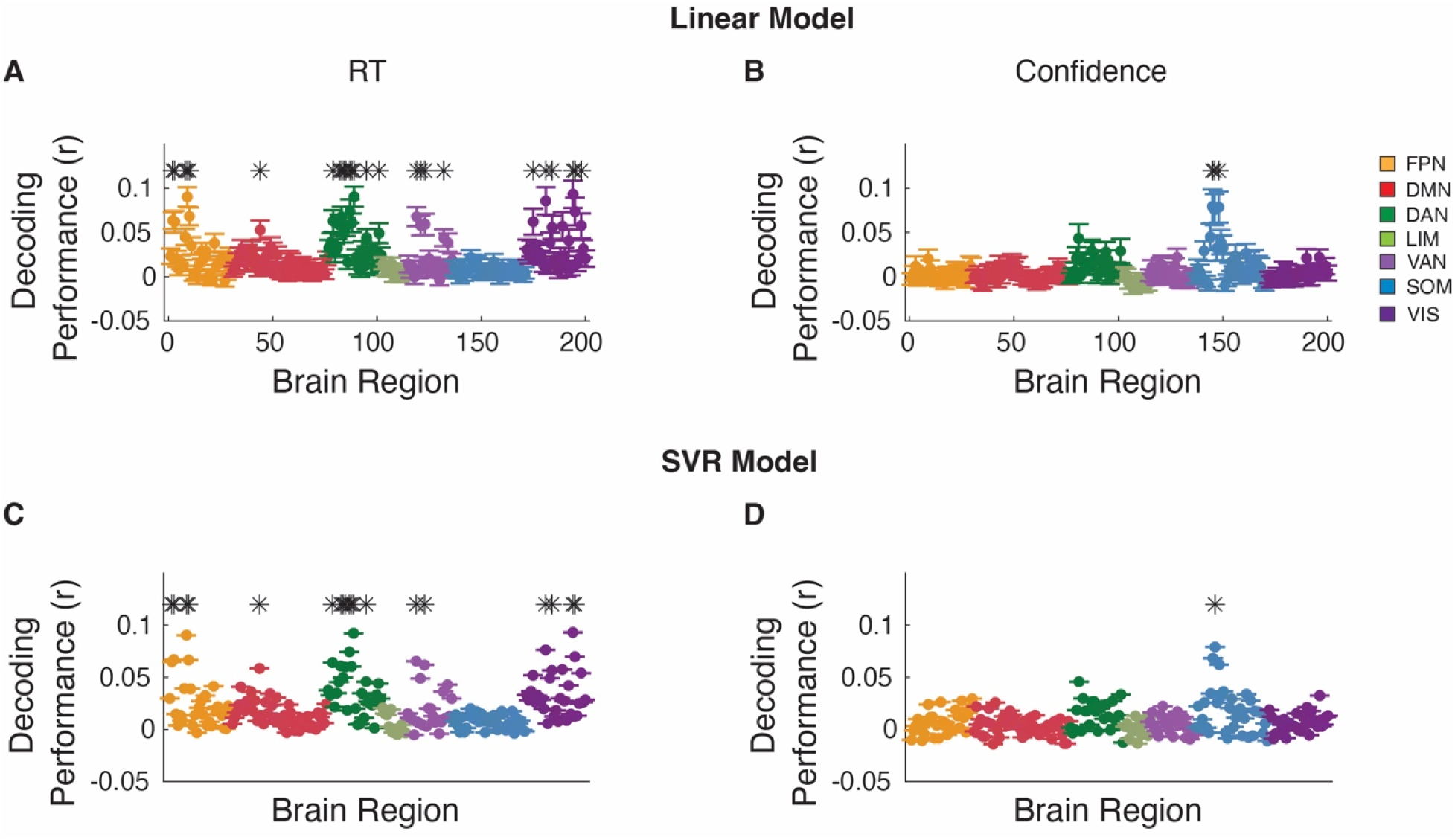
Group-level RT and confidence decoding in Dataset 2. Brain regions from which RT could be decode from using (A) linear and (B) support vector regression (SVR) model at the group level after correcting for 200 comparisons. (C, D) same as plane A and B, but for confidence. * Indicates brain region that significantly, P < 0.05, Bonferroni corrected. FPN, Frontal Parietal Network; DMN, Default Mode Network; DAN, Dorsal Attention Network; LIM, Limbic Network; VAN, Ventral Attention Network; SOM, Somatomotor Network; VIS, Visual Network.

**Figure 8.**
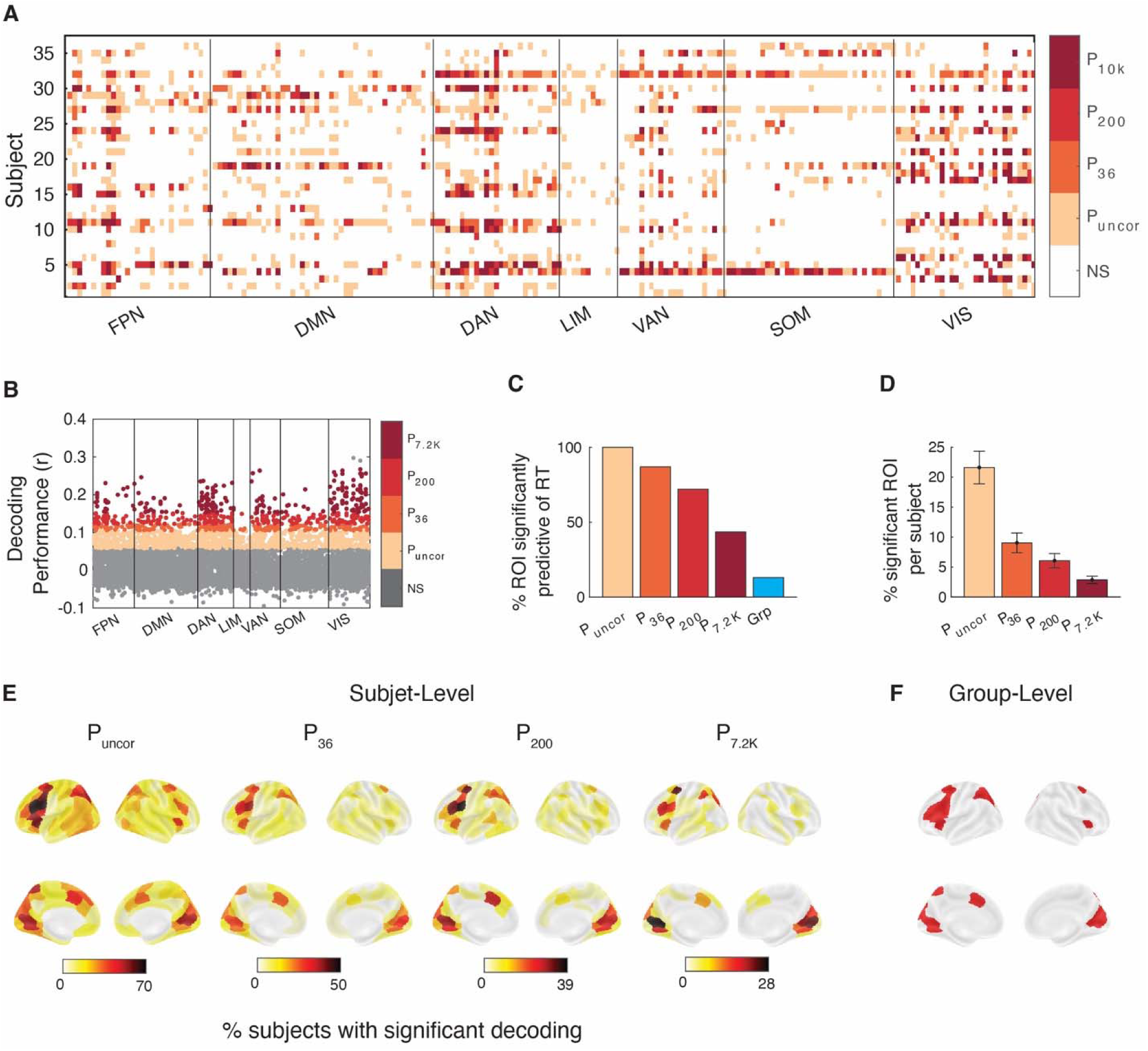
RT can be decoded from across the brain in Dataset 2. (A) Brain regions from which RT could be significantly decoded in each subject and region of interest. Significance is shown at four levels of correction: without any correction (P_uncor_), and after correcting for 36 (P_36_), 200 (P_200_), and 7,200 (P_7.2K_) comparisons. NS, not significant. Colors indicate the most conservative threshold at which one can significantly decode RT from a given region (see color legend on the right). (B) Decoding performance for which subject and brain region. Performance was estimated using the Pearson correlation between empirical and predicted RT. (C) Percentage of ROIs where RT could be significantly decoded for at least one subject for four levels of multiple comparison correction. Group-level results have been added for comparison. (D) Percentage of ROIs per subject where RT could be significantly decoded at four levels of multiple comparison correction. (E) Brain maps plotting percentage of subjects for whom RT could be significantly decoded for four levels of multiple comparison correction. (F) Brain maps plotting ROIs for which RT could be significantly decoded at the group-level after multiple comparison correction. FPN, Frontal Parietal Network; DMN, Default Mode Network; DAN, Dorsal Attention Network; LIM, Limbic Network; VAN, Ventral Attention Network; SOM, Somatomotor Network; VIS, Visual Network.

**Figure 9.**
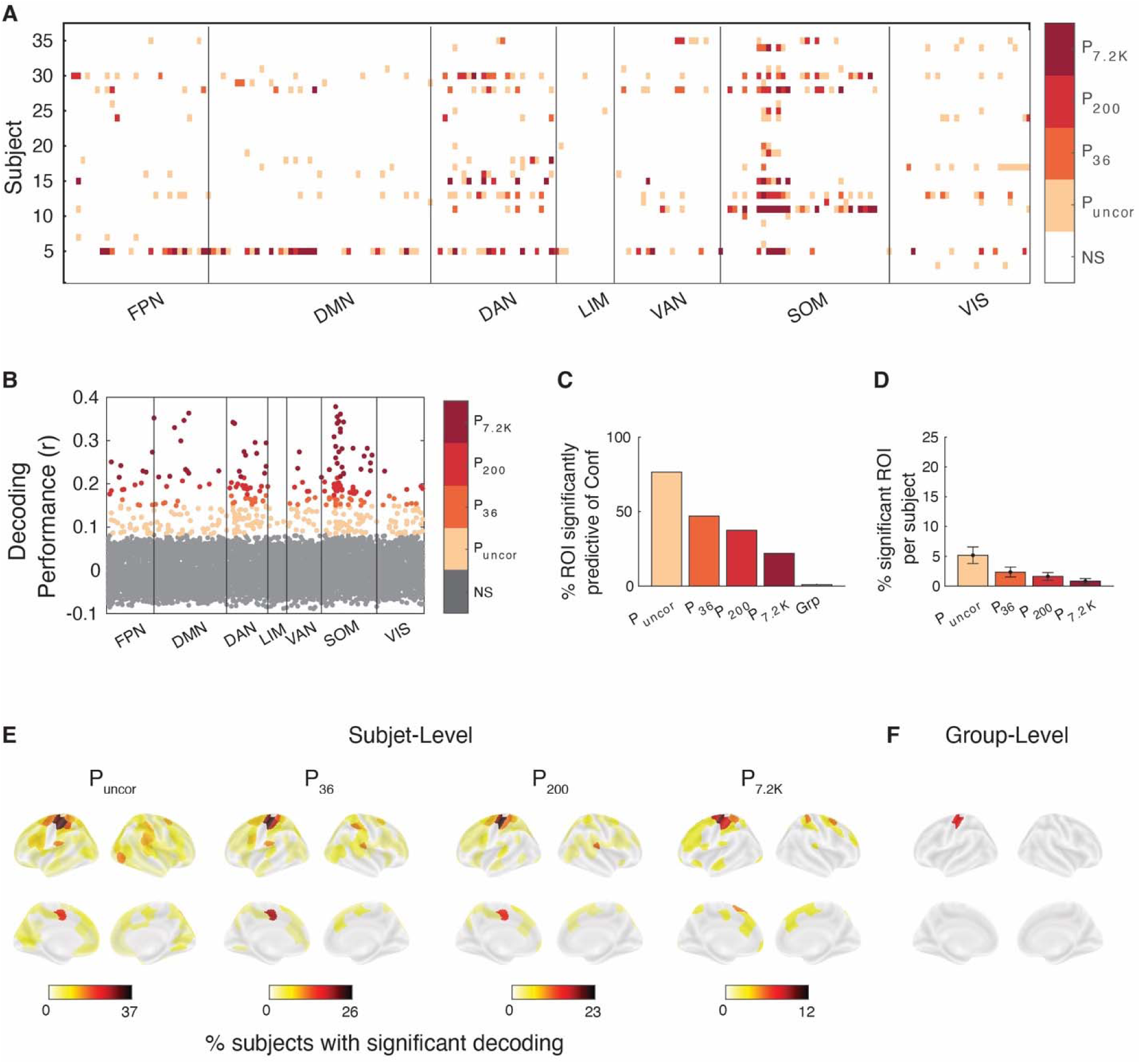
Confidence can be decoded from across the brain in Dataset 2. (A) Brain regions from which confidence could be significantly decoded in each subject and region of interest. Significance is shown at four levels of correction: without any correction (P_uncor_), and after correcting for 36 (P_36_), 200 (P_200_), and 7,200 (P_7.2K_) comparisons. NS, not significant. Colors indicate the most conservative threshold at which one can significantly decode confidence from a given region (see color legend on the right). (B) Decoding performance for which subject and brain region. Performance was estimated using the Pearson correlation between empirical and predicted confidence. (C) Percentage of ROIs where confidence could be significantly decoded for at least one subject for four levels of multiple comparison correction. Group-level results have been added for comparison. (D) Percentage of ROIs per subject where confidence could be significantly decoded at four levels of multiple comparison correction. (E) Brain maps plotting percentage of subjects for whom confidence could be significantly decoded for four levels of multiple comparison correction. (F) Brain maps plotting ROIs for which confidence could be significantly decoded at the group-level after multiple comparison correction. FPN, Frontal Parietal Network; DMN, Default Mode Network; DAN, Dorsal Attention Network; LIM, Limbic Network; VAN, Ventral Attention Network; SOM, Somatomotor Network; VIS, Visual Network.

### Opposite brain-behavior relationship among subjects for the same brain region in Dataset 2

In Dataset 1, we found that for many ROIs, the relationship between brain activity and both RT and confidence often went in the opposite direction for different subjects. Here we show that the same effects also occur in Dataset 2. Similar to Dataset 1, we found that without multiple comparison corrections, high brain activity predicted higher RT in at least one subject and lower RT in at least one subject in 50.0% of all ROIs and 16.5, 6.5, 0.5% after correcting for 36, 200 and 7,200 comparisons, respectively (**Fig. 10A-D**). Reflecting the lower number of trials per subject, confidence showed weaker effects with 12.0% of all ROIs showing significant decoding in different direction for at least two subjects, and 3.0, 1.5, and 0% after correcting for 36, 200, and 7,200 comparisons, respectively (**Fig. 10E-H**). Interestingly, differences emerged between Datasets 1 and 2 when comparing which network contained the most regions with opposite relationship between brain activity. These results demonstrate that the relationship between brain activity and behavioral outcomes is not universal, and that this opposite relationship is not limited to a specific set of regions but instead depends on the individual subject.

**Figure 10.**
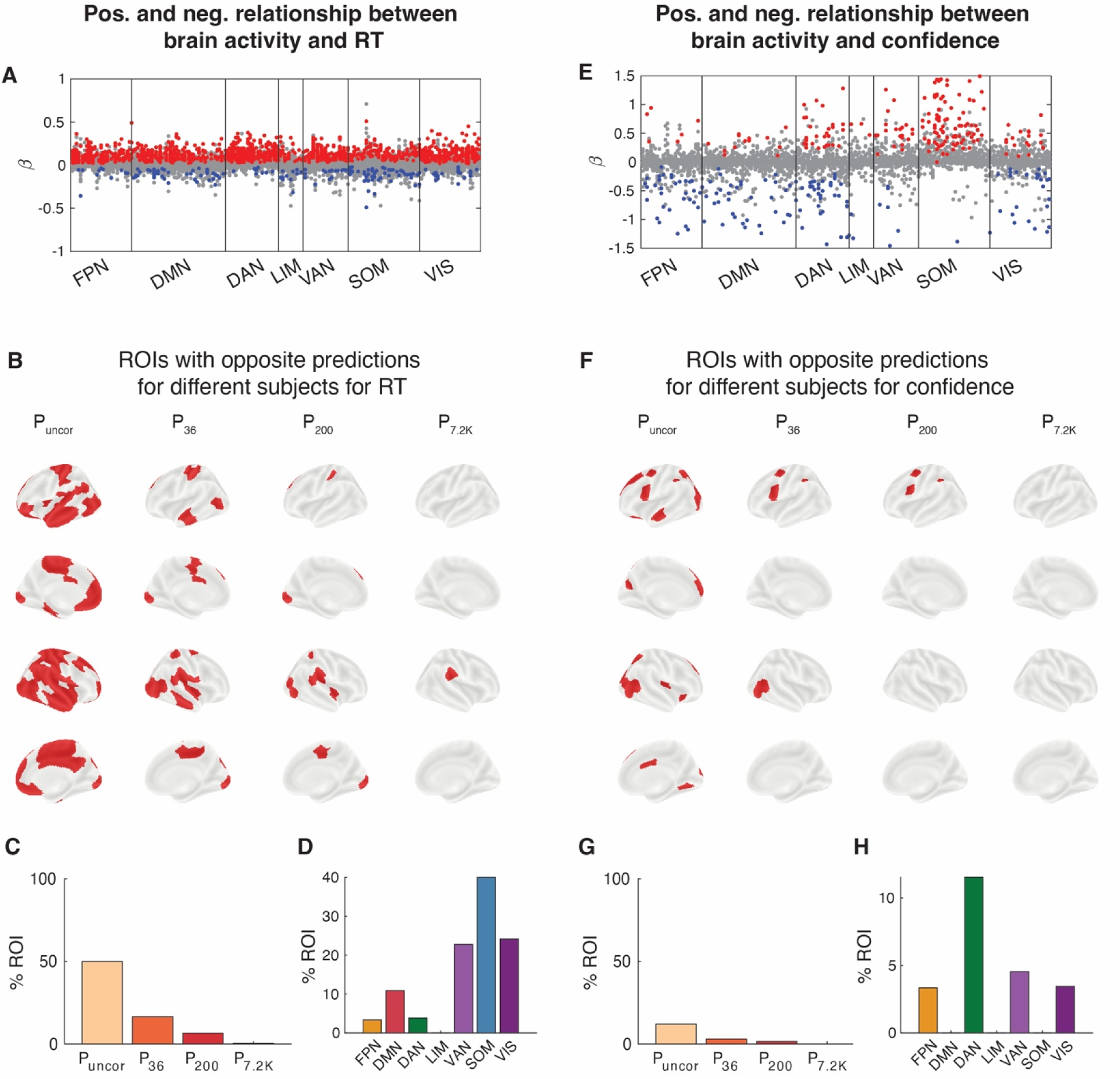
Opposite relationship between brain activity and both RT and confidence among subjects for the same brain region in Dataset 2. (A) Beta values for each subject and region of interest. Red and blue dots reflect subjects for whom RT could be significantly decoded from a given ROI (P < 0.05, uncorrected for multiple comparison). Gray dots show subjects for whom RT could not be significantly decoded from a given ROI. (B) Brain maps showing the ROIs that contained at least one subject with both positive and negative beta values for RT. Significance is shown at four levels of correction: without any correction (P_uncor_), and after correcting for 36 (P_36_), 200 (P_200_), and 7,200 (P_7.2K_) comparisons. (C) Percentage of ROIs that contained at least one subject with both positive and negative beta values for RT for each for four levels of multiple comparison correction. (D) Percentage of ROIs within each brain network that that contained at least one subject with both positive and negative beta values for RT after correcting for 50 comparisons. (E-H) Same as Panels A-D but for confidence.

### No relationship between decoding performance and frame displacement

To confirm that these results were not due to motion artifacts, we determined if there was an association between decoding performance and frame displacement (FD). We performed a regression analysis where average FD for a subject was used to predict average decoding performance for that subject. We found no significant association between FD and decoding performance in either Dataset 1 (RT: R^2^ = 0.02, P = 0.34; Conf: R^2^ = 0.002, P = 0.72; **Fig. S2A-B**) or Dataset 2 (RT: R^2^ = 0.05, P = 0.20; Conf: R^2^ = 0.04, P = 0.22; **Fig. S2C-D**). These results suggest that decoding performance was not driven by motion artifacts.

## Discussion

Traditionally, the brain-behavior relationship has been examined at the group-level to identify the commonalities among individuals. Group-level analyses have typically associated behavioral signatures within a constrained set of brain areas. Here, in contrast to the traditional approach, we focus on the brain-behavior relationship within an individual. We tested how well trial-level RT and confidence can be decoded from the activation in each of the 200 cortical regions of interest obtained using the Schaefer atlas. We showed that RT and confidence can be significantly decoded for at least one subject from brain activity across most of the cortex. Additionally, we were still able to identify differences in the brain-behavior relationship among individuals even with the strictest multiple comparison corrections indicating that these differences were robust and persistent across individuals. These results demonstrate that behavior can be predicted from a wider set of brain areas than would be suggested by standard group analyses.

These findings indicate that individual variability is a major factor in the ability to decode behavioral signatures. In fact, we found that most brain regions could predict RT and confidence in only a small percentage of subjects. Therefore, individual-level analyses might be more sensitive since group-level analyses aggregate over the large degree of within-subject variability (Fisher et al., 2018; Lebreton et al., 2019).

How meaningful is it to find that an ROI predicts behavior in only a small percentage of subjects? For example, one may only be interested in finding brain regions for which behavior can be decoded in most participants. However, the question whether a brain area can predict behavior in at least one subject vs. whether it does so in the majority of subjects are simply different. The latter is concerned with what is common across subjects. This is traditionally the goal of the majority of cognitive neuroscience. While this is an important goal, here we focus specifically on individual differences. In other words, we want to ask how some subjects may differ in meaningful ways from the group. To answer this type of question, one cannot use the traditional approach of looking at the group data. Instead, the idea is that robust decoding in an ROI for even one subject is something that is interesting and meaningful, even if the ROI cannot be used for decoding in any other subject.

A natural question that might arise is whether these individual differences would disappear if the analysis used subject-specific parcellation (Kong et al., 2021). As observed in Kong et al., using subject-specific parcellation can improve brain-behavior decoding. However, in their study, subject-specific parcellation only marginally improved group-level decoding suggesting that the low decoding values do not simply reflect parcellation differences between subjects.

Individual variability in brain-behavior relationship may have many sources. Such variation could emerge from functional degeneracy, the ability of different brain regions to perform the same computation (Price and Friston, 2002; Sajid et al., 2020; Tononi et al., 1999). Functional degeneracy could arise from differences in structural connectivity among individuals (Bansal et al., 2018; Muldoon et al., 2016), since structural connectivity has been associated with differences in behavioral performance (Kanai and Rees, 2011).

Despite the heterogeneity among individuals, a few brain regions showed relatively high consistency across subjects. The lateral frontal cortex and the visual cortex were the most consistent regions from which RT could be decoded. Similarly, for confidence, the somatomotor system was the most consistent area across subjects from which confidence could be decoded, suggesting an association between action-related brain signals and confidence (Fleming et al., 2015; Kiani and Shadlen, 2009).

In a seminal paper, Marek et al. (2022) showed that thousands of subjects are required to robustly estimate the brain-behavior relationship. Even if a brain region can be used to decode behavioral performance, the underlying relationship may not be consistent across subjects underscoring the role of individual differences and the limitation of group-level analyses. Our results suggest that beyond the number of subjects, an arguably even more important factor for brain-behavior relationships is the high degree of between-subject variability. Specifically, if one brain-region is predictive of behavior in one direction in some people but in the opposite direction in others, then no matter how many people a model is trained on, it will always fail to capture many of the individual subjects. It has been suggested that beta values cannot be interpreted particularly from non-linear decoding models using multiple voxels because the beta-values may indicate reduction in noise (Kriegeskorte et al., 2008). This may well be true in some cases, but in the cases where opposite beta values are meaningfully different between individuals, individual variability becomes a crucial limitation in our understanding of how behavior arises from brain activity. Specifically, most brain-behavior computational models assume that there is a uniform relationship between a given region and behavior across all individuals, but this may simply not be true. Instead, we may need to build models that are more flexible and can account for opposite beta values for different subjects.

Finally, we developed a simple test that future studies can use to determine the optimal number of trials in the training and testing bins. This is a crucial step in all decoding analyses, but one that has received little attention. The test relies on training the model on a subset of trials that ranges from 5 to 95% of all trials, with testing performed on the remaining trials. In the context of the current study, we found that several hundred (∼300) trials per person are needed to robustly decode the brain-behavior relationship at the individual level. These results suggest that many previous studies estimating brain-behavior relationship at the individual level may be underpowered. We suggest that the optimal number of trials used in training a decoding model could be based on: (1) minimizing the variance in the decoding performance; or (2) maximizing the difference between decoding performance and the variance in the performance. In the current study, we opted to minimize the variance in the decoding performance even though it came at the cost of a lower decoding performance.

In conclusion, our findings show that behavioral signatures can be decoded from a much broader range of cortical areas than previously recognized. These results highlight that studying the brain-behavior relationship should be studied at both the group and the individual level.

## Author Contributions

J.N. and D.R. designed analysis; J.Y. and D.R. designed study; J.Y., J-H.K., and S-P.K. acquired data; J.N. and J.Y. pre-processed data; J.N. analyzed the data; J.N. and D.R. wrote first draft of paper.

## Acknowledgements

This work was supported by the National Institute of Health (award: R01MH119189) and the Office of Naval Research (award: N00014-20-1-2622).

## Supplemental Information

**Figure S1.**
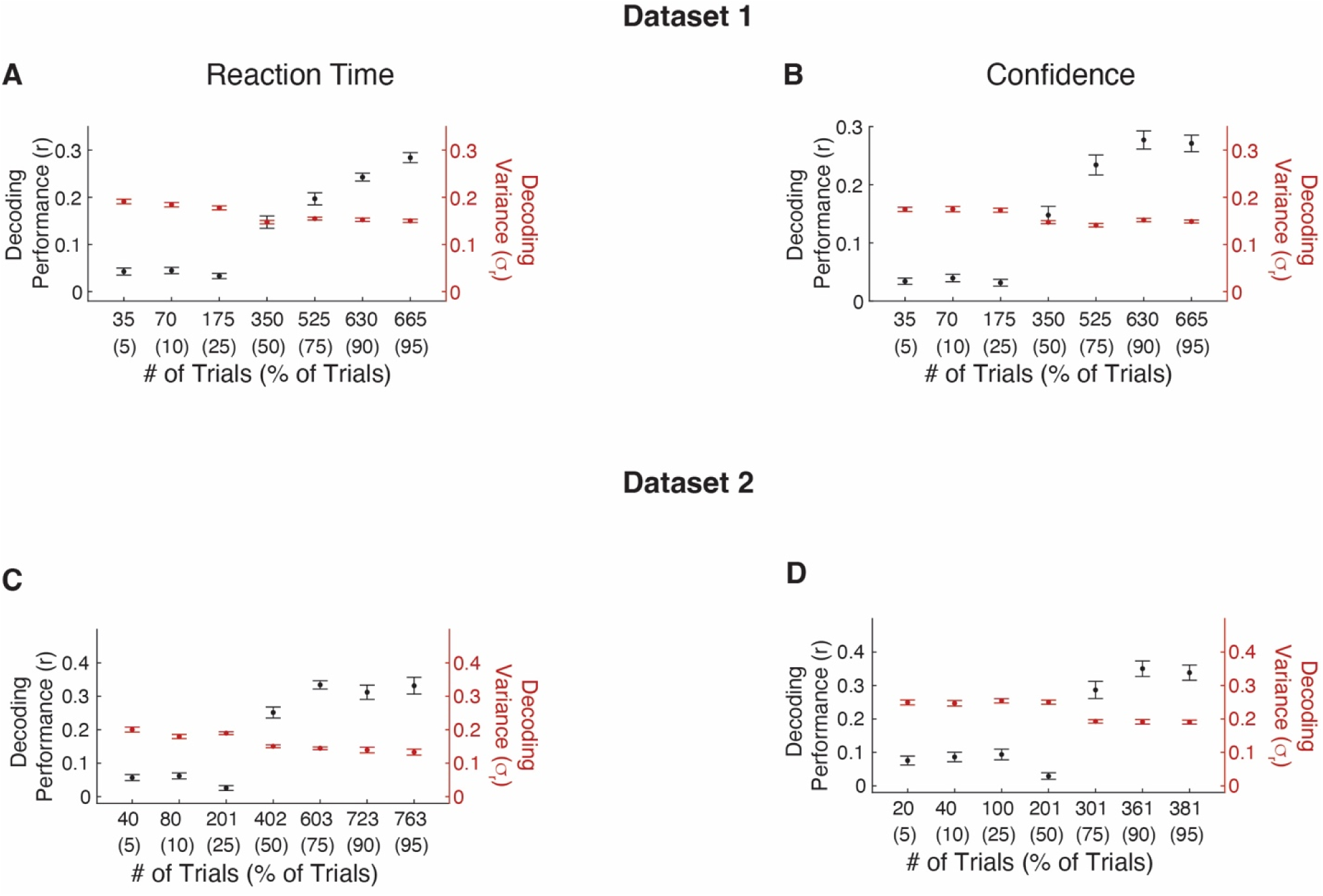
Robust decoding of RT and confidence requires several hundred trials when the number of trials in the test bin is fixed. In Dataset 1, the variance in decoding performance was minimized when using 350 trials (corresponding to 50% of all trials) for both (A) RT and (B) confidence. The black dots present the average decoding performance across subject after 25 repeats for each subject (*left axis)*. The red dots present the average subject-level decoding variance across the 25 repeats for each subject (*right axis)*. The number of test trial is fixed to 5% of the total trials. Error bars show SEM. (C-D) Same as Panels A-B but for Dataset 2. Note that in Dataset 2, confidence was measured on only have the trials (402) and correspondingly the decoding variance was minimized for a higher percentage of trials (75%).

**Figure S2.**
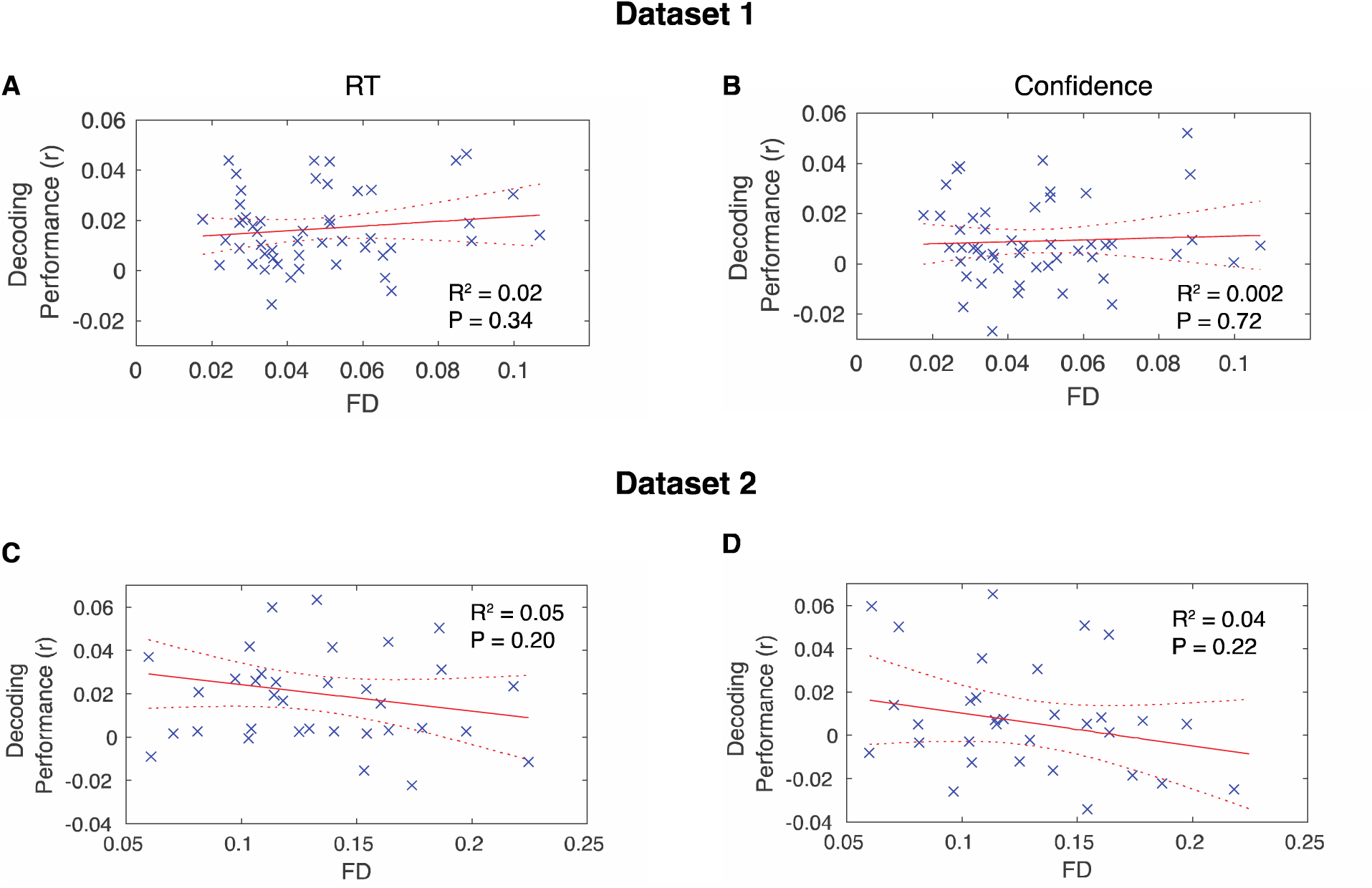
No relationship between decoding performance and frame displacement. No significant (P < 0.05) association exists between decoding performance and frame displacement (FD) in both Dataset 1 (A, B) and Dataset 2 (C, D) for both RT and confidence.

